# Ndc80/Nuf2-like protein KKIP1 connects a stable kinetoplastid outer kinetochore complex to the inner kinetochore and responds to metaphase tension

**DOI:** 10.1101/764829

**Authors:** Lorenzo Brusini, Simon D’Archivio, Jennifer McDonald, Bill Wickstead

**Affiliations:** School of Life Sciences, University of Nottingham, UK

## Abstract

Kinetochores perform an essential role in eukaryotes, coupling chromosomes to the mitotic spindle. In model organisms they are composed of a centromere-proximal inner kinetochore and an outer kinetochore network that binds to microtubules. In spite of its universal function, the composition of kinetochores in extant eukaryotes can differ greatly, and understanding how these different systems evolved and now function are important questions in cell biology. In trypanosomes and other Kinetoplastida, the kinetochores are extremely divergent, with most components showing no detectable similarity to proteins in other systems. They may also be very different functionally, potentially binding to the spindle directly via the inner kinetochore protein KKT4. However, we do not know the extent of the trypanosome kinetochore and proteins interacting with a highly divergent Ndc80/Nuf2-like protein, KKIP1, suggest the existence of more centromere-distal complexes. Here we use quantitative proteomics from multiple start-points to define a stable 9-protein kinetoplastid outer kinetochore (KOK) complex. Two of these core components were recruited from other nuclear processes, exemplifying the role of moonlighting proteins in kinetochore evolution. The complex is physically and biochemically distinct from KKT proteins, but KKIP1 links the inner and outer sets, with its C-terminus very close to the centromere and N-terminus at the outer kinetochore. Moreover, trypanosome kinetochores exhibit intra-kinetochore movement during metaphase, primarily by elongation of KKIP1, consistent with pulling at the outer kinetochores. Together, these data suggest that the KOK complex, KKIP5 and N-terminus of KKIP1 likely constitute the extent of the trypanosome outer kinetochore and that this assembly binds to the spindle with sufficient strength to stretch the kinetochore.

## Introduction

Kinetochores are complex multi-protein machines that ensure the faithful segregation of eukaryotic chromosomes by coupling them to the mitotic spindle and coordinating their movement. The kinetochores of the most closely-studied eukaryotes consist of two major networks: the inner kinetochore constitutive centromere-associated network (CCAN) of ~16 proteins, and the outer kinetochore KMN network consisting of the Knl1-Mis12-Ndc80 complexes (for reviews see (Musacchio and Desai, 2017; Nagpal and Fukagawa, 2016; Varma and Salmon, 2012)). These major networks approximately correspond to the inner and outer plates of electron-opaque material seen at kinetochores by electron microscopy (Pesenti et al., 2016). Microtubules attach to the outer plate. This interaction is primarily mediated by direct binding to microtubules of the Ndc80 complex (Cheeseman et al., 2006; DeLuca et al., 2006) – a hetero-tetramer of Ndc80/Hec1, Nuf2, Spc25 and Spc24 (Ciferri et al., 2005; Wei et al., 2005; Wigge and Kilmartin, 2001). The Ndc80 complex is highly elongated, with a long axis of 55-60 nm (Huis In ’t Veld et al., 2016; Wei et al., 2005). Microtubule binding activity is mediated by calponin homology (CH) domains near the N-termini of Ndc80 and Nuf2 (Ciferri et al., 2008; Wei et al., 2007), which are positioned at the outer kinetochore, while Spc24 and Spc25 bound to the C-termini of Ndc80/Nuf2 interact with components of the CCAN (Malvezzi et al., 2013; Nishino et al., 2013; Schleiffer et al., 2012), thus indirectly connecting the spindle to the centromeres.

Many CCAN and KMN components are conserved between yeast and animals (Kitagawa and Hieter, 2001; Meraldi et al., 2006; Wigge and Kilmartin, 2001) but homologues can also be found in species widely distributed across eukaryotic diversity and many components can be traced back to the last eukaryotic common ancestor (Meraldi et al., 2006; van Hooff et al., 2017). In spite of this ancient origin for complex kinetochores, extant organisms display substantial differences in the repertoire of kinetochore components they encode (van Hooff et al., 2017) and the evolutionary history of kinetochores appears to be associated with both rapid loss in specific lineages (e.g., in the losses of CCAN components and centromeric histone CenH3 in some insects (Drinnenberg et al., 2014)) and large-scale alteration (e.g. in *Tetrahymena*, where Ndc80 is the only component of either CCAN or KMN networks that can be detected in the predicted proteome (van Hooff et al., 2017)). Such disparity in a complex that is both ancient and essential raises important questions about the evolution of this system and how these very different kinetochores function.

The most extreme examples of architectural dissimilarity in eukaryotic kinetochores currently described are those from Kinetoplastida – a group of protozoa including trypanosome and *Leishmania* species, which are important parasites of humans and other animals. Kinetoplastids lack centromeric histone CenH3/CenpA (Lowell and Cross, 2004). Centromeres are instead constitutively marked by 2 kinases (KKT2 and KKT3) with putative DNA-binding motifs, and KKT4, which binds microtubules both in vitro and in vivo and also DNA in vitro (Akiyoshi and Gull, 2014; Llauró et al., 2018). A further 17 KKT (*k*inetoplastid *k*ine*t*ochore) proteins are recruited to kinetochores in a cell-cycle dependent manner, at least some of which are required for correct chromosome segregation (Akiyoshi and Gull, 2014; Nerusheva and Akiyoshi, 2016) in addition to a previously-identified chromosome passenger complex (CPC) containing an Aurora kinase homologue (Li et al., 2008; Tu et al., 2006). KKT proteins (excepting KKT1 and 20) were identified by iterative immunopurification. Since KKTs possess both DNA-binding and microtubule-binding activity, and are a biochemically self-consistent set, it was possible that they encompassed the full extent of the kinetoplastid kinetochore. However, the identification in trypanosomes of a protein with weak similarity to Ndc80 and Nuf2 that associates with the kinetochore, is positioned distal to the KKTs (i.e. more distance from the centromeres) and is essential for karyokinesis, demonstrated additional essential parts of the trypanosome kinetochore (D’Archivio and Wickstead, 2017). This KKT-interacting protein 1 (KKIP1) does not co-precipitate with most KKTs under common conditions for immunopurification, but shows clear association when complexes are stabilized by limited cross-linking (D’Archivio and Wickstead, 2017). Stabilization of complexes also identified six new kinetochore proteins (KKIP2-7), most of which appear to be more centromere-distal than the KKTs.

The discovery of new kinetochore components is suggestive of a possible island of biochemical stability existing distal to KKT proteins that forms the outer plaque of the kinetoplastid kinetochore. Identifying the composition of such a kinetoplastid outer kinetochore complex, if it exists, is of clear importance to understanding the molecular architecture of these unusual kinetochores, including how the outer and inner kinetochores are linked and what role is played by the outer kinetochore if microtubule binding is mediated by centromere-proximal KKT4. Here, we use quantitative proteomics from multiple start-points to test the extent and composition of the trypanosome outer kinetochore. We then use kinetochore components to address the molecular connection between inner and outer sets and test for evidence of outer kinetochore-microtubule binding *in situ* in cells.

## Results

### Quantitative proteomic analysis of KKIP proteins

Previously, we have used quantitative enrichment of proteins co-purifying with KKIP1 under limited cross-linking to demonstrate interaction with KKT proteins (D’Archivio and Wickstead, 2017). This also identified 6 new kinetochore components, KKIP2-7, that are present at the kinetochore but did not co-purify with KKT proteins under standard conditions (Akiyoshi and Gull, 2014). The majority of these new components are downstream of KKIP1 based on co-dependency and localisation analyses, suggesting that they are part of a more centromere-distal set (D’Archivio and Wickstead, 2017), but the extent of this distal set and whether it encompasses stable subcomplexes is not currently known. To test the extent and composition of potential additional complexes at the trypanosome kinetochore, we tagged each of KKIP2-7 in insect-form *Trypanosoma brucei* by integration of coding sequence for YFP at the N-terminus of the endogenous genes and affinity purified the tagged protein (without cross-linking). Co-purifying proteins were then identified by tandem mass spectrometry and relative amounts estimated using label-free normalized spectral index quantification (Trudgian et al., 2011). Spectral intensities and enrichment data for non-redundant trypanosome proteins detected in these experiments are included in Supplemental Data File 1 and original data are available via ProteomeXchange with identifier PXD015100. Note that we do not observe YFP-KKIP4 distributed along the spindle in addition to kinetochores (D’Archivio and Wickstead, 2017) as seen for KKIP4-3HA in procyclic cells (Zhou et al., 2018), suggesting that either position of the tag or modification of the 3’-UTR (as a result of C-terminal tagging) for this gene alters bulk localisation.

Use of semi-quantitative proteomic methods allows for comparison of specific protein enrichment from different purifications. Patterns of co-purifying proteins were assessed by principal component analysis of the relative abundance of proteins identified by immunoprecipitation of KKIP2-7 or in our previous experiments with KKIP1 (D’Archivio and Wickstead, 2017). The first 2 principal components encompass 70% of the total variance in the data and clearly show clustering of KKIP2, 3, 4 and 6 with respect to co-purifying proteins (Fig. 1). KKIP5 and KKIP7 cluster differently consistent with their different temporal localisation (KKIP5 is rapidly lost at anaphase onset and KKIP7 is specifically enriched at metaphase kinetochores (D’Archivio and Wickstead, 2017; Zhou et al., 2019)), but both show clear enrichment of other KKIP components over controls (Suppl. Fig. S1) – suggesting more transient interactions with a core complex. A number of co-purifying proteins are also near neighbours of the KKIP2-4,6 set in this analysis, including common contaminants of immunopurification (e.g. α/β-tubulin) but also several proteins of unknown function/localisation that are enriched over controls in multiple purifications (Suppl. Fig. S1).

**Figure 1.**
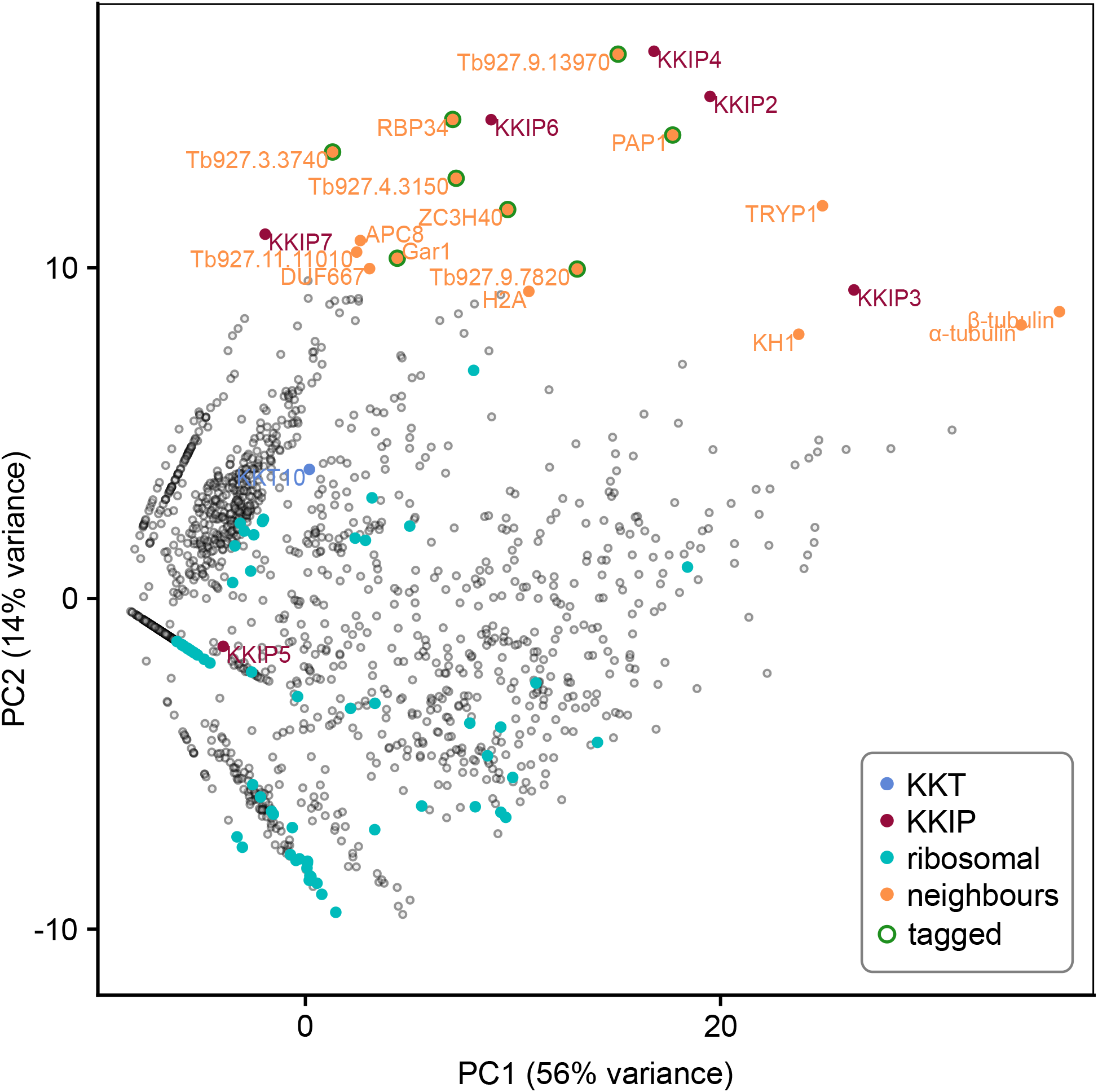
Identification of novel KKIP-interacting proteins. First and second principal components (PC) of integrated spectral intensities for proteins identified by label-free semi-quantitative mass spectrometry as co-purifying with YFP-labelled KKIP1-7 (without crosslinking). Previously identified kinetochore proteins (KKT and KKIP) and near-neighbours to the KKIP2,3,4,6 cluster are highlighted. Ribosomal proteins are highlighted as examples of high-abundance negative controls. Eight close neighbours tagged in this study are also indicated.

### New components suggest a link between kinetochore function and RNA processing

To identify potential new kinetochore components, proteins with biochemical profiles in immunopurification similar to the KKIP2-4,6 cluster were identified. A total of 16 near-neighbours were considered (Fig. 1). From this set, we excluded proteins with demonstrated non-kinetochore localisation. This excludes proteins KH1 and α/β-tubulin, as well as APC8 and histone H2A (all of which may interact with the kinetochore, but predominantly localise elsewhere), plus a tryparedoxin peroxidase TRYP1 (Tb927.9.5770) which is a common contaminant in immunopurification under these conditions. The remaining 8 proteins were tagged by insertion of *YFP* at the N-terminal end of the endogenous coding sequence, with correct integration of the tag being confirmed by Western blotting (Suppl. Fig. S2).

In agreement with their position in principal component analysis, 5 of 8 tagged near-neighbours of KKIP2-4,6 have a clear kinetochore localisation by native fluorescence microscopy (Fig. 2A, B). Following on from existing names, these are referred to herein as KKIP8 to 12. Similarly to KKIP2-6 and most KKTs, newly identified components KKIP9-11 possess no domains of known function in current Pfam profiles. However, KKIP8 (Tb927.3.3160) and KKIP12 (Tb927.11.3340) have known or predicted roles in RNA processing. KKIP8 is one of 2 canonical poly(A) polymerases in trypanosomes, PAP1 and PAP2. PAP1 depletion has no detectable effect on mRNA polyadenylation, but causes an elevation of long non-coding RNA levels and precursors of small nucleolar RNAs (Chikne et al., 2017; Koch et al., 2016) KKIP12/RBP34 is a predicted RNA-binding protein that interacts with the trypanosomal homologue of Mkt1p in yeast 2-hybrid screens (Singh et al., 2014).

**Figure 2.**
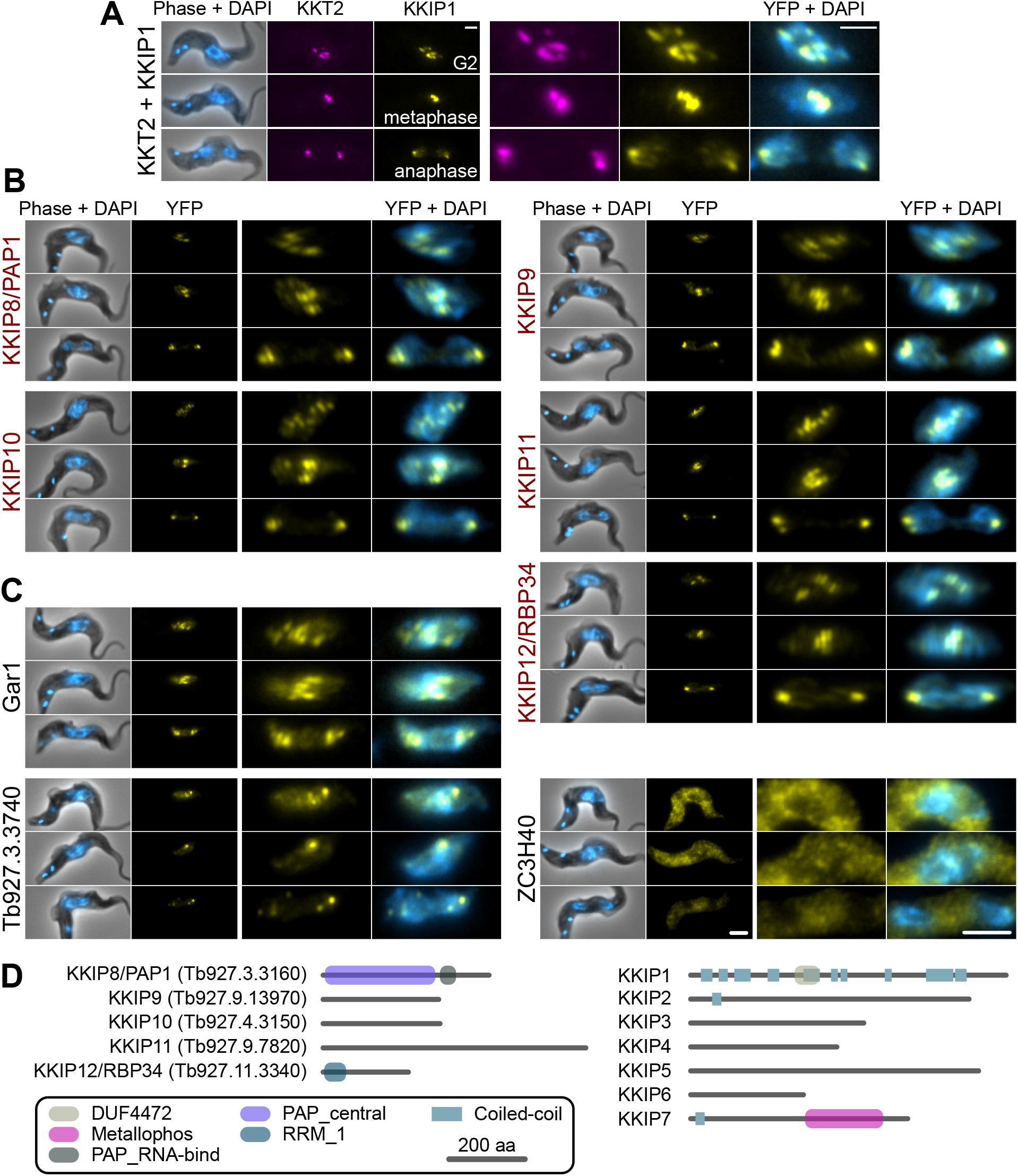
KKIP-interacting proteins include new kinetochore components. Micrographs of native fluorescence in bloodstream-form trypanosomes expressing either known kinetochore components KKT2 and KKIP1 (A), newly identified KKIPs (B), or other proteins detected as co-purifying with KKIPs (C). All proteins are tagged at their N-termini and fluorescence from mScarlet-I (magenta) or YFP (yellow) is shown. Counterstaining of DNA with 4′,6-diamidino-2-phenylindole (DAPI; cyan) and phase-contrast images are also shown. Representative images from cells in G2, metaphase and anaphase are shown for each cell line. Scale bar: 2 μm. (D) Predicted protein architectures for new (KKIP8-12) and previously identified (KKIP1-7) KKT-interacting proteins. Pfam domains with expectation values ≤ 10^−3^ and possible regions of coiled-coil (ncoils; p ≥ 0.5, minimum length 8, window size 21) are highlighted.

All of the newly identified components exhibit temporal patterns of kinetochore binding similar to KKIP1-3 and KKIP6, forming distinct foci in cells from S-phase onwards and being present throughout mitosis (Fig. 2, Suppl. Fig. S3 and (D’Archivio and Wickstead, 2017)). In addition to these components, near-neighbour Gar1 (Tb927.2.3160) localises to mitotic kinetochores but is also present in additional foci present in interphase nuclei (Fig 2C). These interphase foci do not co-localise with KKT proteins that constitutively bind centromeres (Suppl. Fig. S4), suggesting that Gar1 is transiently recruited to kinetochores only during division. In contrast, 2 predicted zinc finger domain-containing proteins – Tb927.3.3740 and ZC3H40 (Tb927.10.14950), showed no clear enrichment at kinetochores and likely represent false positives.

### KKIP8-12 are outer kinetochore components

We have previously demonstrated that loading of KKIP2, 3 and 5 to kinetochores is downstream of KKIP1, reflecting the localisation of these proteins to a position in the kinetochore centromere distal to KKT components (D’Archivio and Wickstead, 2017). Given their behaviour in immunopurification experiments, it is predicted that newly identified KKIPs would be outer kinetochore components. To test this, we generated a cell line in which a marker of the inner (KKT2) and outer (KKIP3) kinetochore were tagged at their endogenous loci with fluorescent markers (mScarlet-I and mTurquoise2, respectively) and used this to determine the position of YFP-tagged KKIP8-12 within the trypanosome kinetochore. Metaphase foci formed by inner and outer kinetochore components can often be distinguished in individual trypanosome cells (for example, (D’Archivio and Wickstead, 2017; Llauró et al., 2018)). In metaphase cells, each of the new kinetochore components KKIP8-12 co-localise with the outer kinetochore component KKIP3 and are distinct from inner kinetochore KKT2 (Fig. 3), providing strong evidence for the majority of the KKIPs being part of physically distinct outer kinetochore complex(es) that do not co-purify with the KKTs of the inner complex.

**Figure 3.**
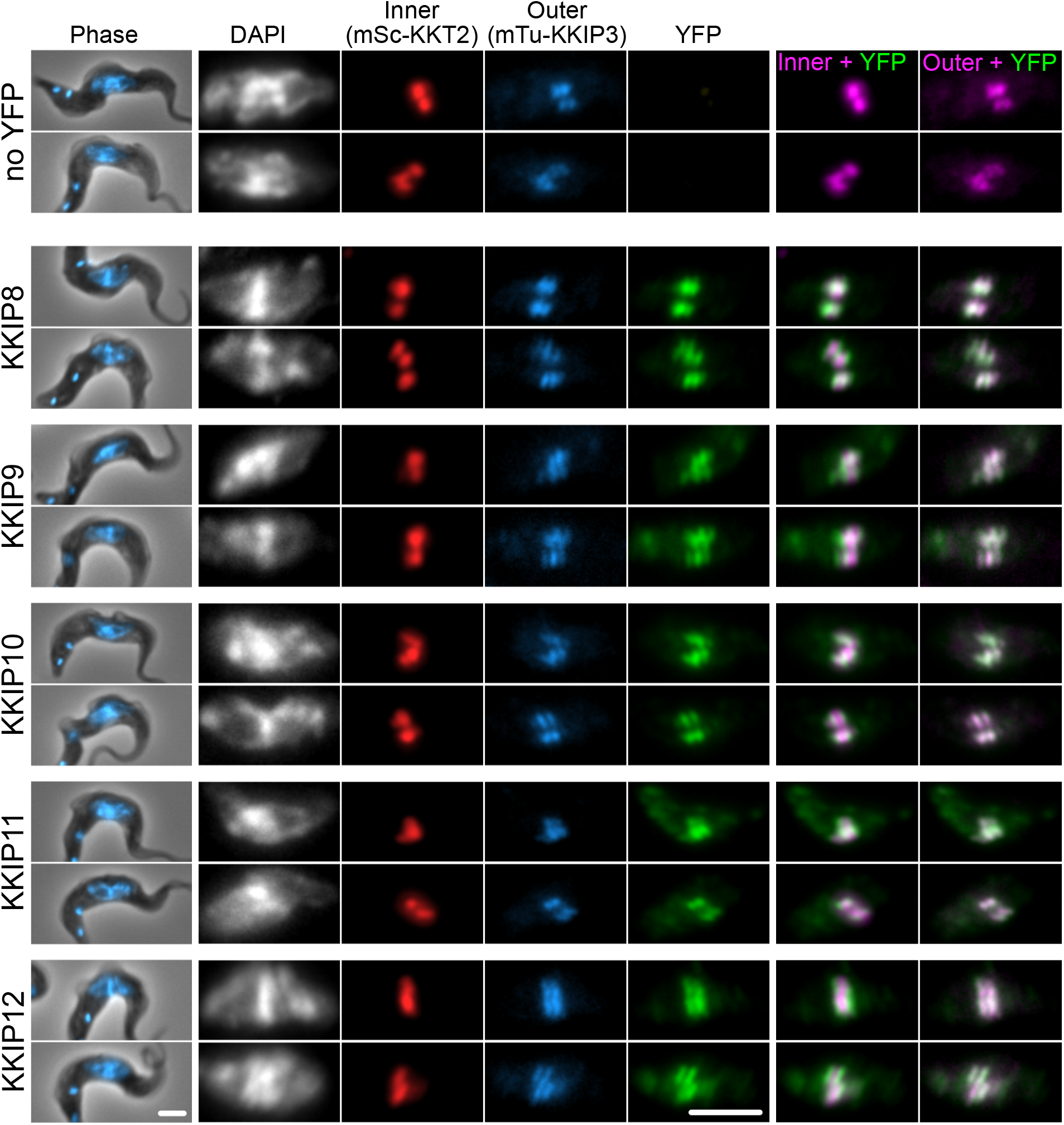
KKIP8-12 are novel outer kinetochore proteins. Micrographs of metaphase bloodstream-form trypanosomes expressing YFP-tagged KKIP8-12 plus inner kinetochore marker KKT2 tagged with mScarlet-I (mSc-KKT2) and outer kinetochore marker KKIP3 tagged with mTurquoise2 (mTu-KKIP3). Counter staining of DNA with DAPI is also shown. Scale bar: 2 μm.

### A stable kinetoplastid outer kinetochore complex

Newly identified KKIPs proteins co-localise to the outer kinetochore. However, it was not known if they are present as one or more subcomplexes. To test for stable (sub)complexes within the KKIP set, and to test for potential further new components, newly identified KKIPs were immunopurified from trypanosomes by the same method as KKIP2-7. Principal component analysis of the normalised spectral intensities for co-purifying proteins clearly shows a distinct group comprising 9 KKIPs (KKIP2-4,6,8-12; Fig. 4A). The separation of these proteins from contaminating hits is substantially improved against immunopurification of only a subset of KKIPs (see Fig. 1) and no additional potential components were identified, suggesting that this is the full extent of the set purifying under these conditions. In addition to non-kinetochore proteins shown above to co-purify with some KKIP proteins (Gar1, ZC3H40 and Tb927.3.3740), near neighbours include spindle components (α/β-tubulin) and proteins that may interact transiently with kinetochores (APC8 and histone H2A). However, TRYP1, KH1, α/β-tubulin and histone H2A are common contaminants in immunopurifications for non-kinetochore nuclear proteins and analysis of enrichment of proteins in KKIP2-12 pull-downs against those co-purifying with a control (YFP-KKIP1) produces a very similar clustering of KKIPs without these likely contaminants (Suppl. Fig. S5).

**Figure 4.**
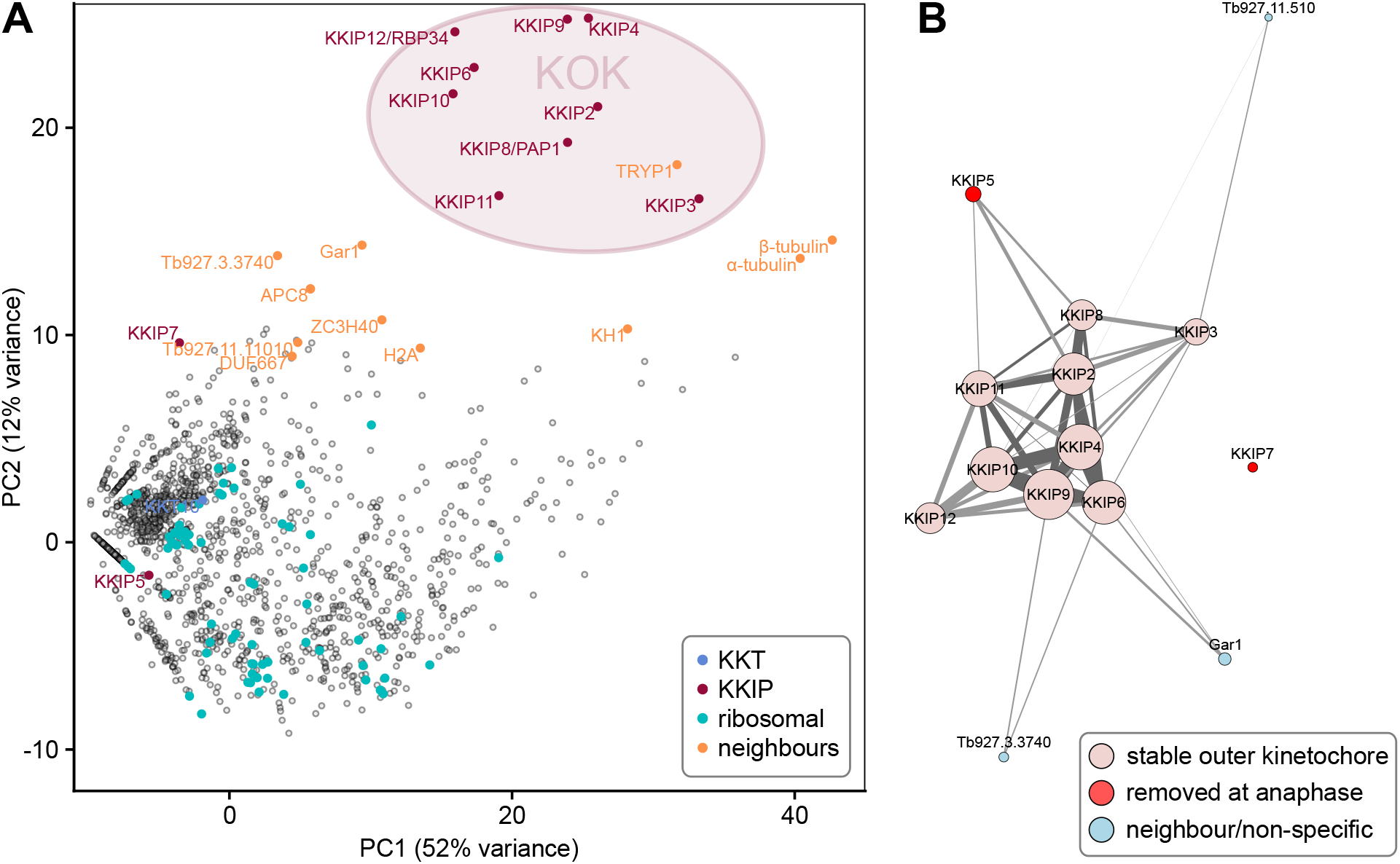
A stable, co-purifying kinetoplastid outer kinetochore complex. (A) First and second principal components (PC) of integrated spectral intensities for proteins identified by label-free semi-quantitative mass spectrometry as co-purifying with YFP-labelled KKIP2-11. A cluster formed by KKIP2-4,6,8-12 defines the extent of a biochemically stable kinetoplastid outer kinetochore (KOK) complex. KKT, KKIP and ribosomal proteins plus near-neighbours from Fig. 1 are highlighted. (B) Network analysis of immunopurification enrichment. To remove likely contaminants, data were processed as log-transformed signal enrichment against non-cross linked KKIP1 immunoprecipitation. All proteins ≥ 8-fold enriched in more than one experiment are shown. Vertex areas are scaled according to sum log enrichment across all experiments. Edge thickness reflects log enrichment of that specific interaction over threshold. Light grey and dark grey edges represent uni- and bi-directional hits, respectively. Vertices are coloured according to the behaviour in the legend. Supplemental Figure S6 shows a similar representation of the network calculated by raw signal.

Network analysis of the most enriched proteins in immunopurifications clearly shows KKIP2-4, 6, and 8-12 form a single coherent complex under these conditions with stable interactions present in multiple experiments (Fig. 4B and Suppl. Fig. S6). These networks also demonstrate the connection of transiently-binding KKIP5 and Gar1 to this core complex. Together with the above, these data strongly suggest the existence of a biochemically stable complex that forms the kinetoplastid outer kinetochore (KOK). From the quantitative purification, KKIP2-4, 6, and 8-12 most likely represent the entirety of the stable KOK, with at least two proteins (KKIP5 and Gar1) being additionally loaded during some stages of mitosis. Significantly, KKT4 (Tb927.8.3680), which has been proposed to be the point of interaction between trypanosome kinetochores and the microtubule (Llauró et al., 2018), is not part of this complex and was not detected in any of the outer kinetochore immunopurifications. This is consistent with the major focus of KKT4 signal being at the inner kinetochore (Llauró et al., 2018) and forming a complex with KKT2 and KKT3 (Akiyoshi and Gull, 2014), implying that either the trypanosome outer kinetochore does not bind to the spindle microtubules, or that this binding is KKT4-independent.

### KKIP1 spans the inner and outer kinetochores and changes length during mitosis

The (inner kinetochore) KKT subcomplexes and KOK complex are biochemically distinct sets. Moreover, the KOK complex does not include microtubule-binding KKT4. This raises two questions: 1) what molecules connect the inner and outer kinetochores in trypanosomes, and 2) does the kinetochore show evidence of grip between the outer complex and the spindle? The first protein localised to the outer kinetochores in trypanosomes was the highly-divergent Ndc80/Nuf2-like protein KKIP1. Stabilization of connections at the kinetochore by limited cross-linking demonstrated that KKIP1 interacts with both KKTs and KKIPs, and that it is required for the localisation of other KKIPs to the kinetochore (D’Archivio and Wickstead, 2017). If KKIP1 acts similarly to Ndc80/Nuf2 in model organisms, it is expected to bridge the inner and outer kinetochores with its C-terminus towards the centromere. To assess the position and orientation of KKIP1 relative to the inner kinetochore and KOK complex, we tagged KKIP1 with YFP at either its N- or C-terminus in cells also expressing KKT2 and KKIP3 tagged with mScarlet-I and mTurquoise2, respectively (see Fig. 3). The manually-assigned positions of foci from each fluorophore were then used to probe the nanometer-scale architecture of the kinetochores by calculating the positions of KKIP1 or KKIP3 in the image plane relative to the closest focus of KKT2 signal (i.e. label separation or ‘delta’; (Joglekar et al., 2009; Wan et al., 2009)). Only cells with clear in-focus KKT2 foci were considered, resulting in sub-pixel position measurements for 315, 392 and 430 foci (mScarlet-I, YFP and mTurquoise2, respectively) from 111 mitotic cells. These measurements exclude any contribution to distances from displacement of either kinetochore or spindle in z. The potential for swivel/k-tilt is discussed below, but the contribution of displacement of the entire spindle will be small as mitotic trypanosomes settle onto glass slides such that the spindle axis lies predominantly in the xy plane (meaning an underestimate of < 3% for a typical 4-μm anaphase spindle with both poles in the focal plane). As spindles are oriented randomly in the xy plane and both ends of the spindle are considered, the contribution of chromatic aberration is expected to sum to zero across all foci/cells.

As seen previously (D’Archivio and Wickstead, 2017), both the N-terminal end of KKIP1 and KKIP3 are significantly displaced distally from KKT2 in anaphase cells (i.e. further towards the poles along the spindle axis; Fig. 5A, B). However, improved imaging of many more mitotic cells shows the distance between the inner and outer domains of anaphase kinetochores is substantially greater than previously estimated (Fig. 5C; mean absolute distance of 76 ± 8 nm and 109 ± 9 nm along the spindle axis for fluorophores at the N-termini of KKIP1 and KKIP3, respectively). In contrast, the position of YFP placed at the C-terminus of KKIP1 is nearly indistinguishable from KKT2 at both anaphase and metaphase kinetochores (Fig. 5B, C; mean distances of 24 ± 9 nm and 14 ± 8 nm respectively). KKIP1 position and orientation are very reminiscent of the Ndc80 complex: highly elongated, with C-terminus binding the inner kinetochore and N-terminus at the outer kinetochore, consistent with the suggestion that KKIP1 is a divergent Ndc80/Nuf2 homologue (D’Archivio and Wickstead, 2017; Llauró et al., 2018)

**Figure 5.**
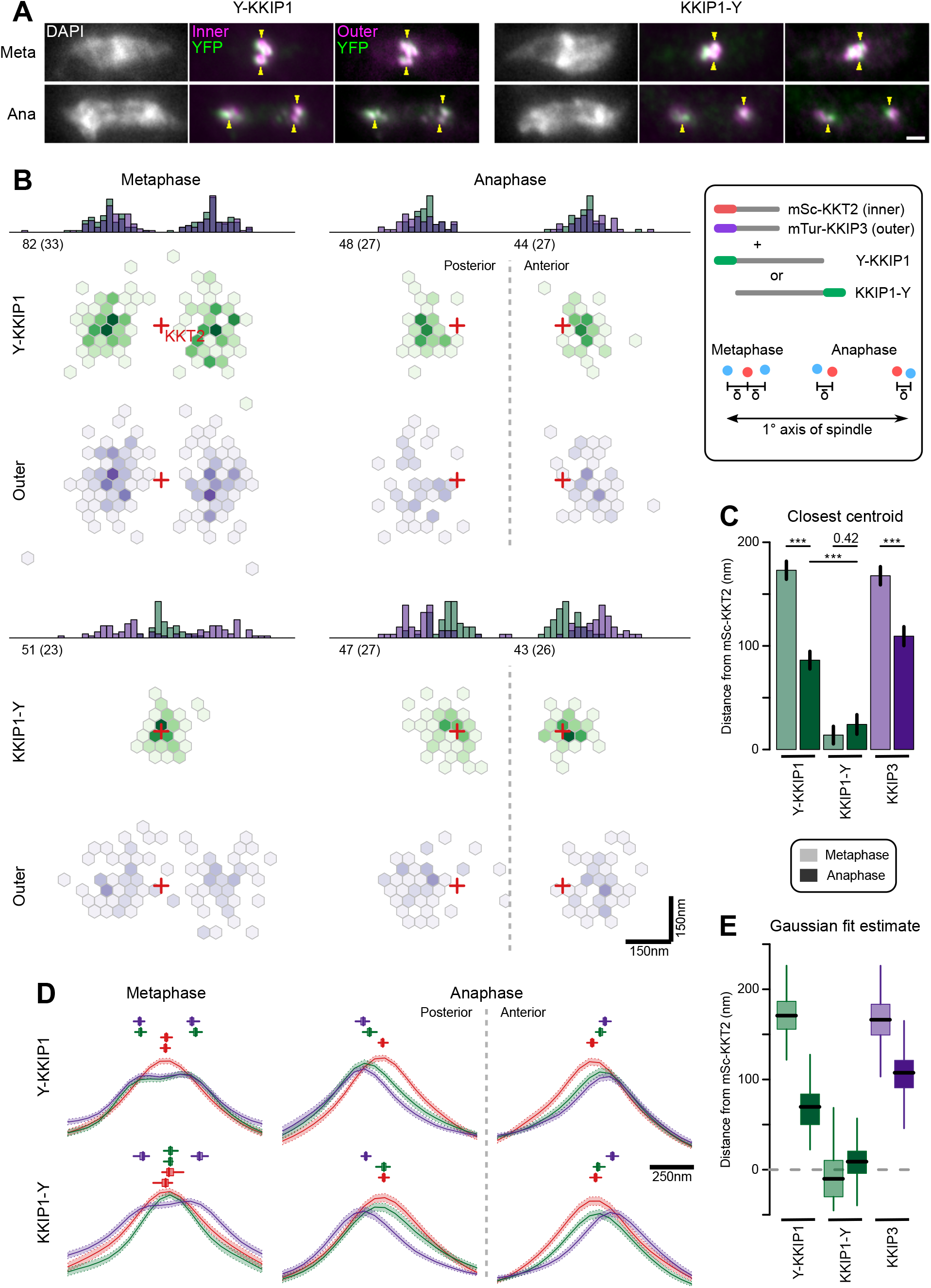
KKIP1 bridges the inner and outer kinetochore and responds to metaphase tension. (A) Representative micrographs of cells expressing either YFP-KKIP1 or KKIP1-YFP and markers for the inner (mSc-KKT2) or outer (mTu-KKIP3) kinetochore. Counterstaining of DNA with DAPI is also shown. For reference, centres of some mSc-KKT2 foci are indicated by arrows. Scale bar: 2 μm. (B) Distribution of centroids of either YFP-KKIP1 or KKIP1-YFP and mTu-KKIP3 relative to the closest centroid of mSc-KKT2. All data are transformed such that the primary spindle axis is horizontal with the posterior end of the cell to the left and are grouped according to stage in mitosis (metaphase or anaphase) and whether YFP-KKIP1 or KKIP1-YFP expressing cells. Red crosses indicate the positions of mSc-KKT2 centroids. Histograms show the 1D distributions along the spindle axis. Numbers indicate total number of foci and independent cells (brackets) captured for each class. Scale is shown bottom-left. (C) Mean distance along the primary spindle of KKIP1 and KKIP3 centroids relative to KKT2. Bars show s.e.m. (*** = p-value < 0.001; Student’s t-test). (D) Fluorescence along the spindle axis (‘line scans’) around mSc-KKT2 foci. Mean (line) and s.e.m. (shaded area) for each channel is shown. Boxes over line scans show centres of either one (anaphase) or two (metaphase) Gaussian distributions fitted to 100 bootstrap replicates of the data (median, interquartile distance and range are shown by bar, box and line, respectively). (E) Distance along the primary spindle of Gaussian-fit estimates for KKIP1 and KKIP3 positions. Bar, box and lines represent median, interquartile distance and ranges for 100 bootstrap replicates of the datasets.

In yeast and animal cells, binding of sister outer kinetochores to opposing sides of the spindle at metaphase is associated with changes in the distance between inner and outer domains along the K-K axis that is reduced either at anaphase or if tension is removed (Joglekar et al., 2009; Maresca and Salmon, 2009; Smith et al., 2016; Uchida et al., 2009; Wan et al., 2009). We hypothesized that if trypanosome outer kinetochores bind to microtubules, they should also show evidence of structural change associated with metaphase tension, whereas binding predominantly via inner kinetochore KKT4 should not induce such change. Consistent with intra-kinetochore stretch, at metaphase the distance along the spindle axis from the outer kinetochore to the nearest KKT2 focus is approximately doubled (173 ± 10 nm and 168 ± 9 nm for YFP-KKIP1 and KKIP3, respectively) relative to anaphase distances (Fig. 5B, C). However, as the foci in metaphase cells represent sister kinetochore pairs (unlike those in anaphase) and the inner kinetochore pairs in trypanosomes do not separate beyond the resolution limit of light microscopy, the observed change in KKT2-KKIP1/3 centroid distances could also be produced by an increase in the inter-sister distance, so long as the inner-inner distances were still insufficient to create two distinguishable foci of mSc-KKT2. A separation of the inner-inner pairs of ~200 nm would entirely account for the observed change in KKT2-KKIP1/3 distance between metaphase and anaphase without any change in the inner-outer kinetochore distance – although it would be inconsistent with the close apposition of bioriented trypanosome kinetochores seen previously by electron microscopy (Ogbadoyi et al., 2000).

To discriminate between intra-kinetochore and inter-sister stretch, we analysed the distribution of signal intensity along the spindle axis in all 3 fluorescence channels around the 315 mSc-KKT2 foci (Fig. 5D). As expected for roughly point-sources, the signal intensities around anaphase foci are approximately Gaussian and the clear poleward displacement for KKIP3 and the N-terminus of KKIP1 seen previously is still evident. Fitting these data to unconstrained Gaussian distributions recapitulates the distance estimates from manually-assigned centroid positions (Fig. 5D, E; 70 nm and 107 nm for YFP-KKIP1 and KKIP3, respectively) in a manner that is robust to input image and initial parameter selection (estimated from 100 bootstrap replicates of the fit). At metaphase, outer kinetochore markers show a clear bimodal distribution across the kinetochore pair, as seen previously (Fig. 3 and (Llauró et al., 2018)). To allow for distance between sister kinetochores, metaphase distributions were fitted to the sum of two Gaussian distributions. Distribution widths for each wavelength were fixed to values from anaphase cells, such that broadening of signal could only be achieved by displacement of the two centres. Again, the unsupervised Gaussian fit estimates for the distance to the outer kinetochores are very similar to those based on centroids (171 nm and 166 nm for YFP-KKIP1 and KKIP3, respectively). In addition, the best fits to the signal distribution for both mSc-KKT2 and KKIP1-YFP place the centres of their paired Gaussian distributions within 20 nm of each other (Fig. 5E). This is in good agreement with the fact that there is very little broadening of either mSc-KKT2 or KKIP1-YFP signal distribution in metaphase relative to anaphase (in contrast to outer kinetochore signals) as would be seen for large inner-inner distances. Together, these data are incompatible with inter-sister kinetochore displacement making a substantial contribution to the distance between outer and inner kinetochore signals at metaphase, and demonstrate that trypanosome kinetochores substantially alter their overall structures during mitosis – being significantly more elongated during biorientation at metaphase than during poleward movement at anaphase.

## Discussion

### The kinetoplastid outer kinetochore

Using quantitative proteomics we have defined a stable 9-protein kinetoplastid outer kinetochore complex including 5 newly identified kinetochore components. This complex is self-consistent by immunopurfication and most components have a similar temporal localisation through the cell cycle. A tenth outer kinetochore protein, KKIP5, which binds to the kinetochore only until anaphase (D’Archivio and Wickstead, 2017), clearly binds via KOK complex components. Rapid reduction in KKIP5 levels at anaphase are likely via the proteasome, as inhibition of proteasome activity stabilizes KKIP5 levels (Zhou et al., 2019)and this is necessary to see KOK interaction in immunopurifications (data not shown).

The ‘full’ extent of any complex in the cellular milieu is dependent on context and experimental approach. As our model of the trypanosome kinetochore improves, it is very possible that additional components will be discovered – especially if such components bind transiently or form interactions that are not stable under conditions used for immunopurification. However, from the data presented here, the 9 components are highly likely to represent the full extent of the stable KOK and, together with KKIP5 and the N-terminal end of KKIP1, are positioned as far from the inner kinetochore as the most distal ultrastructural features identified for the trypanosome kinetochore (Ogbadoyi et al., 2000). Significantly, none of these outer kinetochore components co-purify with KKT4, which is found at the inner kinetochore, but is the only component to date shown to have an intrinsic microtubule-binding capacity (Llauró et al., 2018).

### Origins of outer kinetochore components

One of the unusual features of kinetoplastid chromosome segregation is the widespread lack of detectable similarity between components of the system and those from models (Akiyoshi and Gull, 2014, 2013; Berriman et al., 2005). The vast majority of proteins identified to date as components of the kinetoplastid kinetochore have no clear homology to proteins outside of the Kinetoplastida (Akiyoshi and Gull, 2014; D’Archivio and Wickstead, 2017) and how this system evolved is an outstanding question. Trypanosomes do possess a functional chromosome passenger complex (Li et al., 2008) and, although it was originally thought the non-kinase components of this were unique to kinetoplastids, there is strong evidence that TbCPC1 is a divergent INCENP orthologue (van Hooff et al., 2017). In addition, the trypanosome outer kinetochore component KKIP1 has weak similarity to Ndc80/Nuf2 family proteins, with which it also shares some functional features, suggesting this and potentially other kinetoplastid kinetochore proteins may be very divergent homologues of more canonical components (D’Archivio and Wickstead, 2017). However, other components – in particular, two pairs of kinases (KKT2,3 and KKT10,19) present in inner kinetochore complexes – clearly demonstrate that at least some parts of the system do not share common ancestry with kinetochores in model systems.

As for most previously identified kinetoplastid kinetochore proteins, 7 of 9 components of the stable KOK have no clear homology to proteins outside of the Kinetoplastida. In contrast, KKIP8 (PAP1) and KKIP12 (RBP34) are clear homologues of proteins involved in RNA binding/processing. What function (if any) these proteins perform at the outer kinetochore is unclear; no detectable segregation defects were observed on RNA-interference-mediated protein knockdown (data not shown). However, both are stable components of the KOK network and their presence demonstrates the divergence of kinetochore composition by incorporation of proteins with other nuclear roles (presumably initially as transiently associating or moonlighting proteins).

### Stretch at the kinetoplastid kinetochore

In model organisms, the structure of the kinetochore changes substantially during mitosis in a manner that is dependent on the mechanical environment. In budding yeast, the separation of centromeric Cse4/CenpA and the N-terminus of Ndc80 along the spindle axis is ~70 nm during metaphase, but reduces by ~25nm in anaphase, most of which is mediated through changes in the conformation of the Ndc80 complex (Joglekar et al., 2009). In human cells, the CenpA/N-Ndc80 distance along the K-K axis is ~100 nm and reduces by ~30 nm if microtubule pulling forces are chemically removed (Dudka et al., 2018; Suzuki et al., 2014; Uchida et al., 2009; Wan et al., 2009). At least some of this reduction in distance is due to displacement of the outer kinetochore away from the K-K axis (i.e. swivel/k-tilt) or other kinetochore distortions (Magidson et al., 2016; Smith et al., 2016) but the Ndc80 complex appears fully extended and aligned to the axis of pull when kinetochores are bioriented at metaphase (Suzuki et al., 2018).

Here we have demonstrated that the unusual kinetochores of trypanosomes also change dimensions during mitosis in a manner consistent with intra-kinetochore stretch. To our knowledge, this is the first such demonstration of structural change at the kinetochore for any species outside of the opisthokonts (the group containing animals and fungi). In trypanosomes this change in intra-kinetochore distance is much more pronounced than for animals/fungi. Allowing for up to 50 nm separation between paired inner kinetochores (consistent with the outer-limits of the bootstrap estimates), the distances between an inner marker (KKT2) and the N-terminus of KKIP1 are ~145 nm at metaphase – around twice their distances during anaphase (Fig. 6). These distance measurement are along the spindle axis only (i.e. Δ_1D_) so are potentially a convolution of stretch and swivel/k-tilt or distortion to the overall structure (Magidson et al., 2016; Smith et al., 2016). Given the compact area of trypanosome kinetochores (each binding 3-5 microtubules), the contribution of distortion is likely to be negligible. Moreover, there is no evidence in the measurements of centroid positions in the xy plane (Fig. 5B) for swivel being a major contributor to reduction Δ_1D_ distances at anaphase.

**Figure 6.**
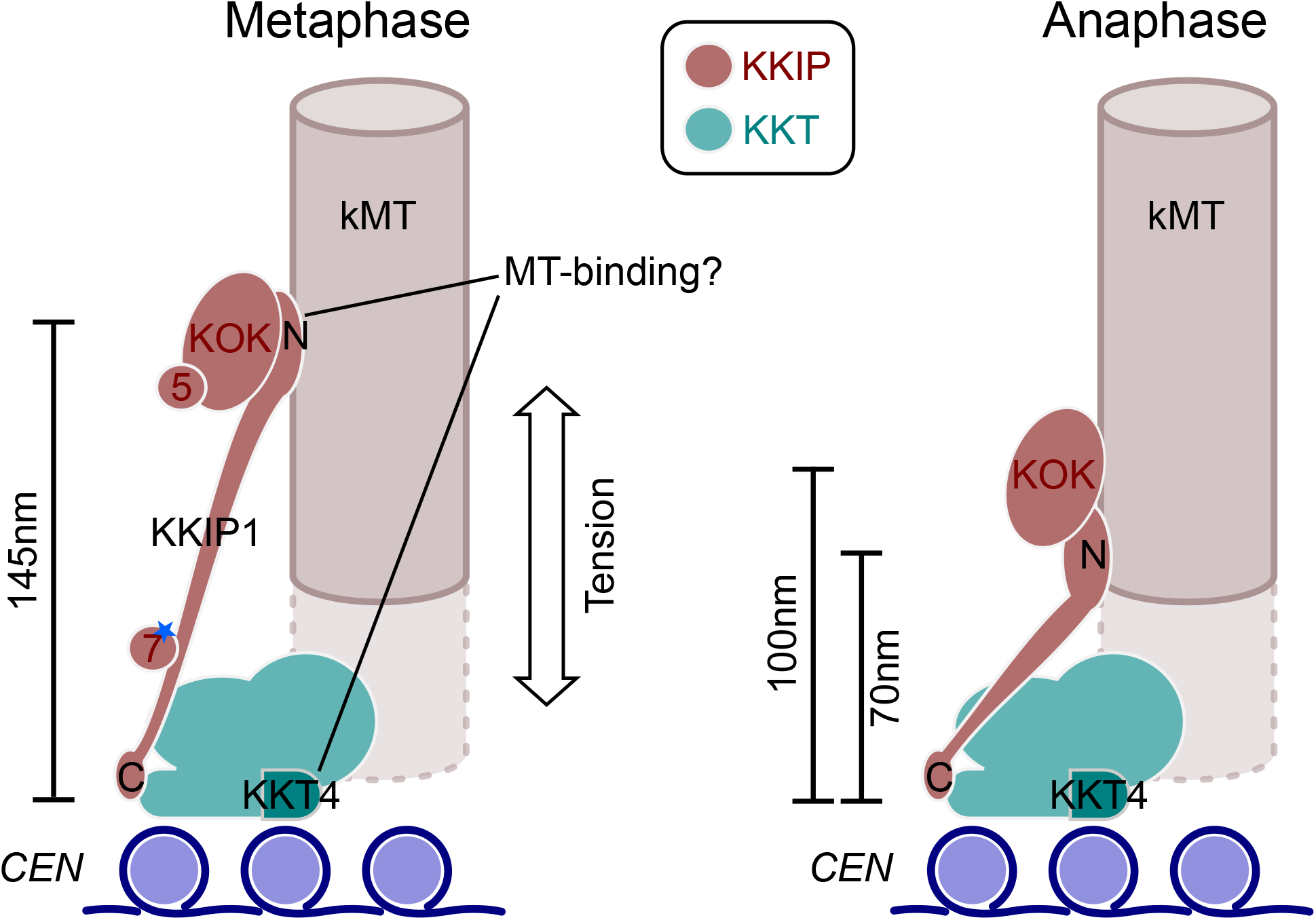
Model for trypanosome kinetochore architecture and alterations between phases of mitosis. Model incorporates data from (Akiyoshi and Gull, 2014; D’Archivio and Wickstead, 2017; Llauró et al., 2018) and work herein.

Most of the change in intra-kinetochore distance in trypanosomes appears to result from changes in the length of KKIP1, which bridges from a position within ~20 nm of KKT2 to the outer kinetochore. Hence, KKIP1 sits in the same orientation as Ndc80, but is much more elongate than even the Ndc80 complex (of which Ndc80 is only part). The structure of KKIP1 is yet to be solved, but its primary sequence is substantially longer than many Ndc80/Nuf2 family proteins and predicted to be dominated by coiled-coils (D’Archivio and Wickstead, 2017) Even so, the maximum extent between N- and C-termini suggests very few globular domains will be present. Aligning the measurements here with previous ultrastructural description of trypanosome kinetochores (Ogbadoyi et al., 2000) places the outer kinetochore not at the outer edge of the ~50 nm electron-dense layer(s) as previously thought, but at the very thin (~10 nm) layer that is seen 40 nm further towards the pole in anaphase cells. We suggest that this electron-dense layer comprises the N-terminal end of KKIP1 and the KOK complex, marking the most distal part of the kinetoplastid kinetochore. Such physical separation likely goes some way to explaining why this section of the kinetochore was not isolated along with the KKTs (Akiyoshi and Gull, 2014; Nerusheva and Akiyoshi, 2016).

The likelihood that kinetoplastid kinetochores respond to metaphase tension raises important questions regarding how attachment is mediated. No proteins encoded in the trypanosome genome contain identifiable Ndc80 or Nuf2 CH domains. KKT4 has demonstrated microtubule-binding capacity, but this protein is immediately centromere-proximal and produces only a mild defect on knockdown compared to other components (Llauró et al., 2018) Evidence of stretch between inner and outer kinetochores is compatible with a microtubule-binding role for inner-kinetochore KKT4 during spindle assembly or kinetochore capture, but make it very unlikely that the outer kinetochore is a passive passenger to metaphase attachments mediated through the inner kinetochore alone. Alternatively, small amounts of KKT4 might be present at the outer kinetochore. KKT4 tagged at either terminus localises to positions on the spindle other than the kinetochores ((Llauró et al., 2018)http://tryptag.org/). However, this additional localisation is mostly at the poles and KKT4 did not co-purify with any KOK component (Fig. 1, 4), nor they with KKT4 (Akiyoshi and Gull, 2014). As such, pulling at the outer kinetochores creating elongation of KKIP1 and movement of the KOK currently appears incompatible with KKT4 being the primary link between kinetochores and the spindle during metaphase and suggests additional/alternative binding from the outer kinetochore.

## Materials and methods

### Cell lines and cell culture

All work was performed using cell lines derived from SmOxP427 and SmOxB427 cells (in the case of procyclic-form or bloodstream-form cells, respectively), which are derivatives of *Trypanosoma brucei brucei* strain Lister 427 modified to express transgenic T7 RNA polymerase and Tet-repressor protein from the tubulin locus (Poon et al., 2012). Procyclic cells were grown at 28°C in SDM79 medium (Brun and Schönenberger, 1979) supplemented with 10% fetal bovine serum. Bloodstream-form cells were grown in HMI-9 medium supplemented with 15% fetal bovine serum at 37°C and 5% CO_2_ (Hirumi and Hirumi, 1989).

All constructs were derived from pEnCY0-H, pEnNY0-H, pEnNmSc0-N or pEnNmTu0-B, which encode YFP and hygromycin resistance marker (pEnCY0-H, pEnNY0-H), mScarlet-I and neomycin resistance marker (pEnNmSc0-N), or mTurquoise2 and blasticidin resistance marker (pEnNmTu0-B). Sequences and maps for these vectors are available at www.wicksteadlab.co.uk. For N-terminal tagging constructs (pEnNxx-x vectors), ~200bp targeting sequences from the N-terminal end of the coding sequence and upstream sequence were ligated downstream of the fluorescent protein coding sequence, along with a linearisation site between the targeting sequences. For the C-terminal tagging construct (pEnCxx-x vectors), ~200bp targeting sequences from the C-terminal end of the coding sequence and downstream sequence were ligated upstream of the fluorescent protein coding sequence, along with a linearisation site between the targeting sequences. All primers used for cloning are available in Supplemental Data File 2. Plasmids were linearized with NotI and transfected into trypanosomes by electroporation as described in (Schumann Burkard et al., 2011). Stable transfectants were selected with 50 μg ml^−1^ hygromycin B, 10 μg ml^−1^ blasticidin or 2.5 μg ml^−1^ G418 in the case of procyclic cells, or 5 μg ml^−1^ hygromycin B, 2 μg ml^−1^ blasticidin or 2.5 μg ml^−1^ phleomycin for bloodstream-form cells. Integration at the endogenous loci was confirmed by diagnostic PCR and Western blotting to confirm expected size of chimeric protein.

### Immuno-purification

Immuno-purification was performed as described in (Daniels et al., 2012) from procyclic form cells. Briefly, ~3×10^9^ cells expressing YFP-tagged kinetochore components were harvested by centrifugation from actively dividing cultures. Cells were washed once in ice-cold HKMEG (150 mM KCl, 150 mM glucose, 25 mM HEPES pH7.8, 4 mM MgCl_2_, 1 mM EGTA) and then with HKMEG containing 5 μM E64-d. Cells were lysed in HKMEG containing 1% (v/v) Nonidet P40, 1 mM dithiothreitol, 20 μM proteasome inhibitor MG-132 and a protease inhibitor cocktail (2 mM 1,10-phenanthroline, 0.5 mM phenylmethanesulfonyl fluoride, 50 μM leupeptin, 7.5 μM pepstatin A, 5 μM E64-d) followed by sonication for 3 min at 30% intensity applied for 30% of the cycle with a Sonopuls 70 W ultrasonic homogenizer (Bandelin). The lysate was then cleared by centrifugation at 20 000×*g* for 30 min. Cleared lysate was allowed to bind for 2 h on ice with gentle agitation to ~5x molar excess of affinity-purified rabbit anti-GFP polyclonal antibodies which had been covalently attached to paramagnetic beads (Dynabeads Protein G, Invitrogen) by dimethyl pimelimidate treatment (Unnikrishnan et al., 2012). Beads were washed extensively in HKMEG containing 0.1% (v/v) Nonidet P40, 0.5 mM dithiothreitol, and bound complex subsequently eluted by incubating beads 3 times in 100 mM glycine pH2.7.

### Mass spectrometry and label-free quantitative analysis

Immuno-purified samples were desalted by precipitation with acetone at −20°C, washed in cold acetone and solubilized in Laemmli sample buffer. Samples were encapsulated in a polyacrylamide matrix by running a short distance into an SDS-PAGE gel, followed by staining with SYPRO Ruby protein stain (Bio-Rad) and excision of gel fragment. Gel fragments were washed with 50% acetonitrile in 50 mM NH_4_HCO_3_ pH8.5, dehydrated in 100% acetonitrile, and air dried. Proteins were digested for 16 h with 20 μg ml^−1^ trypsin (Promega) in 25 mM NH_4_HCO_3_ pH8.5 at 37°C. Mass spectrometry was performed on an LTQ Orbitrap XL mass spectrometer (Thermo Scientific) at the University of Oxford Central Proteomics Facility (www.proteomics.ox.ac.uk).

Label-free quantitation was performed from mzXML data files using the Central Proteomics Facilities Pipeline at the Advanced Proteomics Facility, University of Oxford (www.proteomics.ox.ac.uk). Data were searched with X! Tandem and OMSSA engines against a custom, non-redundant protein database of predicted protein sequences from TREU927/4 strain (www.tritrypdb.org) with the inclusion of exogenous protein sequence and common contaminating peptides. Possible modification of peptides by N-terminal acetylation, carbamidomethylation (C), oxidation (M), and deamidation (N/Q) was permitted in searches. Peptide identifications were validated with PeptideProphet and ProteinProphet (Nesvizhskii et al., 2003) and lists compiled at the peptide and protein level. iProphet was used to combine search engine identifications and refine identifications and probabilities. Normalized spectral index quantitation (SINQ) was applied to the grouped metasearches to give protein-level quantitation between labeled samples and controls, as described in (Trudgian et al., 2011), and implemented by the Central Proteomics Facilities Pipeline at the University of Oxford. SINQ values are summed intensities of matched fragment ions for all spectra assigned to a peptide (identified by ProteinProphet), normalized for differences in protein loading between datasets and for individual protein length. Only proteins with at least 2 detected peptides and an estimated false discovery rate of ≤ 1% relative to a target-decoy database were considered. A total of 1582 distinguishable trypanosome proteins were detected across all experiments. Mass spectrometry proteomics data have been deposited to the ProteomeXchange Consortium via the PRIDE (Perez-Riverol et al., 2019) partner repository with the dataset identifier PXD015100. Processed data are also provided in Supplemental Data File 1.

Enrichment and principal component analysis was performed in the statistical programming package ‘R’ (www.r-project.org). Quantitative values were analysed as either log-transformed SINQ values (for principal component analysis) or log-transformed ratio of sample SINQ value versus KKIP1 control immunopurification (enrichment and network analysis).

### Protein localisation by fluorescence microscopy

For analysis of localisation of tagged proteins by native fluorescence, cells were harvested from mid-log phase cultures, washed twice in phosphate-buffered saline (PBS; 137 mM NaCl, 3 mM KCl, 10 mM Na_2_HPO_4_, 1.8 mM KH_2_PO_4_) and allowed to settle onto glutaraldehyde-derivatized silanized slides. Cells were fixed for 5 min in 2% (w/v) formaldehyde, permeabilised in −20°C methanol for at least 2 min, re-hydrated in PBS and incubated with 15ng ml^−1^ 4′,6-diamidino-2-phenylindole for 5 min, before mounting in 1% (w/v) 1,4-diazabicyclo[2.2.2]octane, 90% v/v glycerol, 50 mM sodium phosphate pH 8.0. All micrographs shown in this manuscript are from bloodstream-form cells, but match localisations of the same proteins tagged procyclic-form cells (data not shown).

Images were captured on an Olympus BX51 microscope equipped with a 100x UPlanApo objective (1.35 NA; Olympus) and Retiga R1 CCD camera (Qimaging) without binning (64.5 nm pixel size). All images of fluorescent proteins were captured at equal exposure settings without prior illumination. Images for level comparison were also processed in parallel with the same alterations to minimum and maximum display levels. Image acquisition was controlled by μManager open source software (Edelstein et al., 2014). Analysis was performed in ImageJ (Schneider et al., 2012) and the statistical programming package ‘R’ (www.r-project.org).

### Analysis of sub-kinetochore localisation

For analysis of relative positions of kinetochore components, cells in mitosis were transformed such that the longest axis of the nucleus (corresponding to the mitotic spindle axis) lay along the x axis in the posterior-anterior direction. Two independent measurements of fluorescence positions were performed. In the first, the sub-pixel peak of signal for individual foci at 3 wavelengths were assigned manually. Peak locations from either YFP-tagged KKIP1 or mTu-KKIP3 were assigned to the closest focus of mSc-KKT2 in xy and relative positions of focus centroids in each channel calculated. In the second, the positions of foci visible from mSc-KKT2 only were recorded. The distributions of fluorescence along the x-axis (‘line scans’) for all 3 wavelengths were then sampled (645 nm either side of the mSc-KKT2 focus). For foci from cells in anaphase, the mean distributions from all sampled foci in each channel at either posterior or anterior end of the spindle were then fitted to single Gaussian distributions by non-linear regression with starting values estimated from the distribution. To allow for signal from both sides of metaphase kinetochores, mean distributions for each channel in metaphase cells were fitted to the sum of two Gaussian distributions of equal peak height and per-channel variance estimated from the measurements for single anaphase foci. To assess robustness and infer confidence intervals, 100 bootstrap datasets were generated by randomly sampling with replacement the foci in each category of cells and fitting Gaussian distributions to the mean distribution as above.

No correction was made for components of the spindle axis in z; elevation of one pole of a typical 4 μm spindle by up to 1 μm in z (sufficient for kinetochore foci to move out of the focal plane) would lead to an underestimate of the distance along the true spindle axis due to only considering xy components by <3%, which is below the precision of the measurements. Full scripts used for transformation and analysis are available from the authors on request.

## Acknowledgements

This work was supported by a BBSRC new investigator award (BB/J01477X/1) and University of Nottingham strategic funding to BW, MRC studentship (1506963) to LB, and BBSRC studentship (1364116) to JM. Mass spectrometry and initial quantitation were performed at the Advanced Proteomics Facility, University of Oxford (www.proteomics.ox.ac.uk). We are grateful in particular for the assistance of Svenja Hester in preparation and running of mass spectrometry samples. We are also grateful to Catarina Gadelha (University of Nottingham) for help with processing and analysis of the semi-quantitative proteomics data following acquisition. Mass spectrometry proteomics data have been deposited to the ProteomeXchange Consortium via the PRIDE partner repository with the dataset identifier PXD015100.

