## Supplemental Figures for "Ndc80/Nuf2-like protein KKIP1 connects a stable kinetoplastid outer kinetochore complex to the inner kinetochore and responds to metaphase tension"

Supplemental Figure S1. Multiple proteins are identified across KKIP immuno-purifications. Label-free semi-quantitative mass spectrometry showing relative enrichment of proteins immunopurified with KKIP1-KKIP7 against KKIP1 under standard, non cross-linking, as well as cross-linking conditions (D'Archivio and Wickstead, 2017). Signals from published proteomic datasets, including kinetochore KKTs and KKIPs, as well as proteins highly enriched in multiple experiments (neighbours) are highlighted. For display, intensity of proteins not detected in a specific immunoprecipitation are set to an arbitrary minimum value.

Supplemental Figure S2. Immunoblots of whole cell lysates from cells expressing YFP- or mStrawberry- (mSt-) endogenously tagged KKIP or KKIP-interacting proteins. Ponceau-S staining of total protein on the membrane is shown as a control for loading.

Supplemental Figure S3. Spatio-temporal localisation of newly identified kinetoplastid kinetochore components through the trypanosome cell cycle. Representative images of cells arranged by inferred position in the cell cycle (G1 to cytokinesis from top to bottom) are shown. Native fluorescence from YFP (yellow), in addition to counterstaining of DNA with DAPI (cyan) and phase contrast images. Scale bars: 2 µm.

Supplemental Figure S4. KKIP-interacting proteins transiently associate with trypanosome kinetochores. Micrographs of bloodstream-form cells at metaphase expressing one of Gar1, Tb927.3.3740 and ZC3H40 endogenously tagged with YFP at the N-terminus. Markers for the inner and outer kinetochore were also tagged with mScarlet-I (KKT2) and mTurquoise2 (KKIP3), respectively. Counterstaining of DNA with DAPI is also shown. Scale bar: 2 µm.

Supplemental Figure S5. KKIP8-KKIP11 interact with the stable kinetoplastid outer kinetochore. Extension of the label-free semi-quantitative mass spectrometry based approach used in Suppl. Fig. 1. Signals from published proteomic datasets, including kinetochore KKTs and KKIPs, as well as common interactors across experiments are highlighted.

Supplemental Figure S6. Network analysis of raw protein levels from immunopurifications. Only proteins occurring in the top 25% of signal in more than one experiment are shown. Vertex areas are scaled according to sum normalised signal across all experiments. Edge thickness reflects normalised signal of interaction over threshold. Light and dark grey edges represent uni- and bi-directional hits, respectively. Vertices are coloured as described in the legend.

D'Archivio S, Wickstead B. 2017. Trypanosome outer kinetochore proteins suggest conservation of chromosome segregation machinery across eukaryotes. *J Cell Biol* **216**:379–391. doi:10.1083/jcb.201608043

Supplemental Figure S1

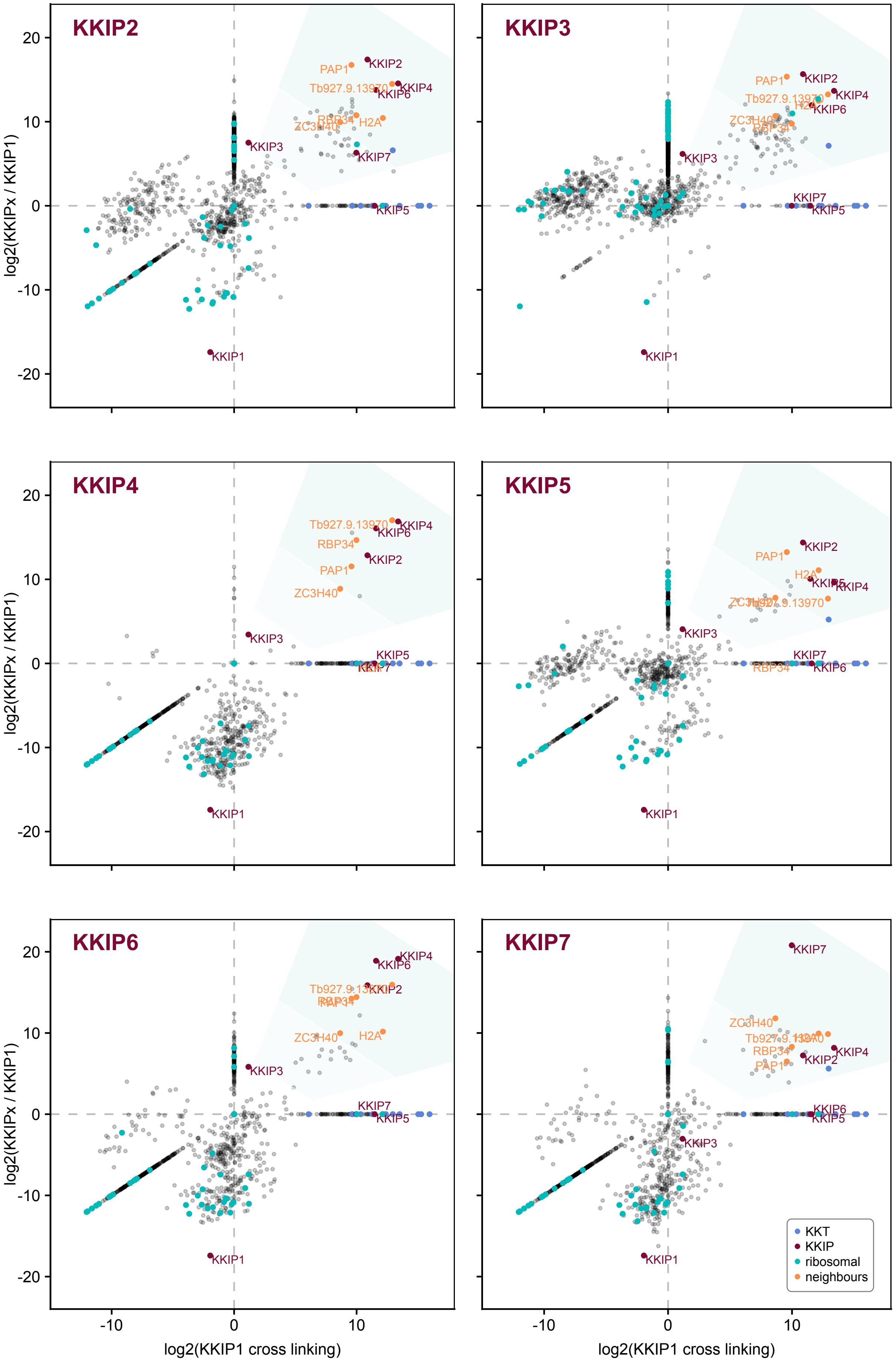

### Supplemental Figure S2

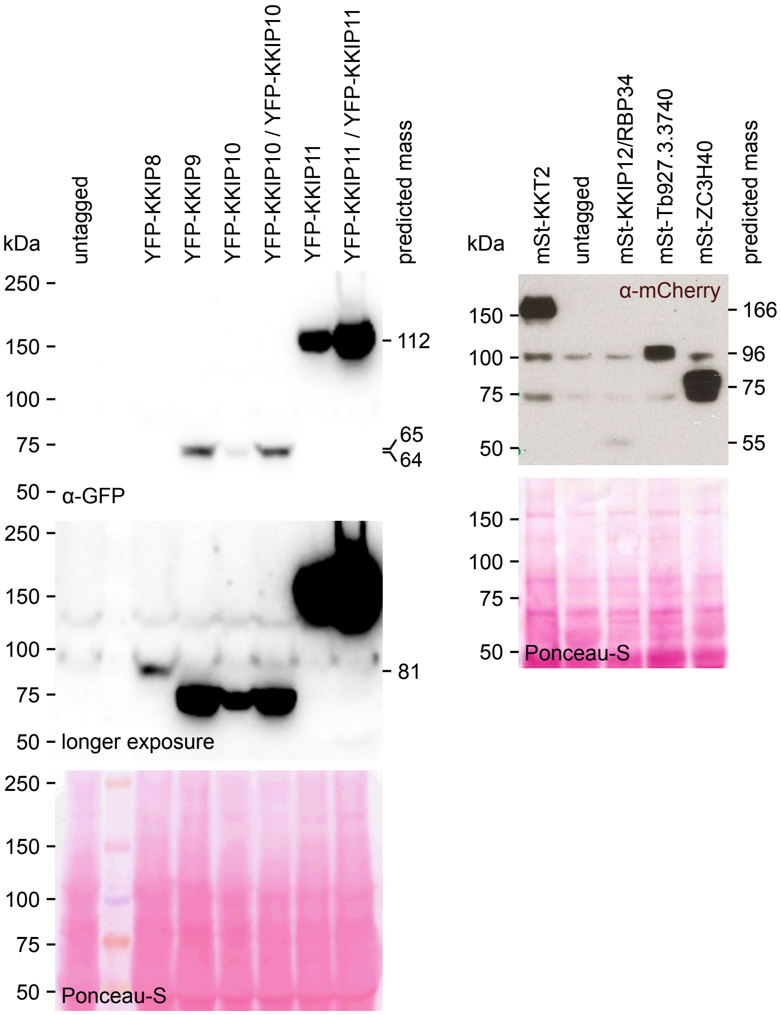

### Supplemental Figure S3

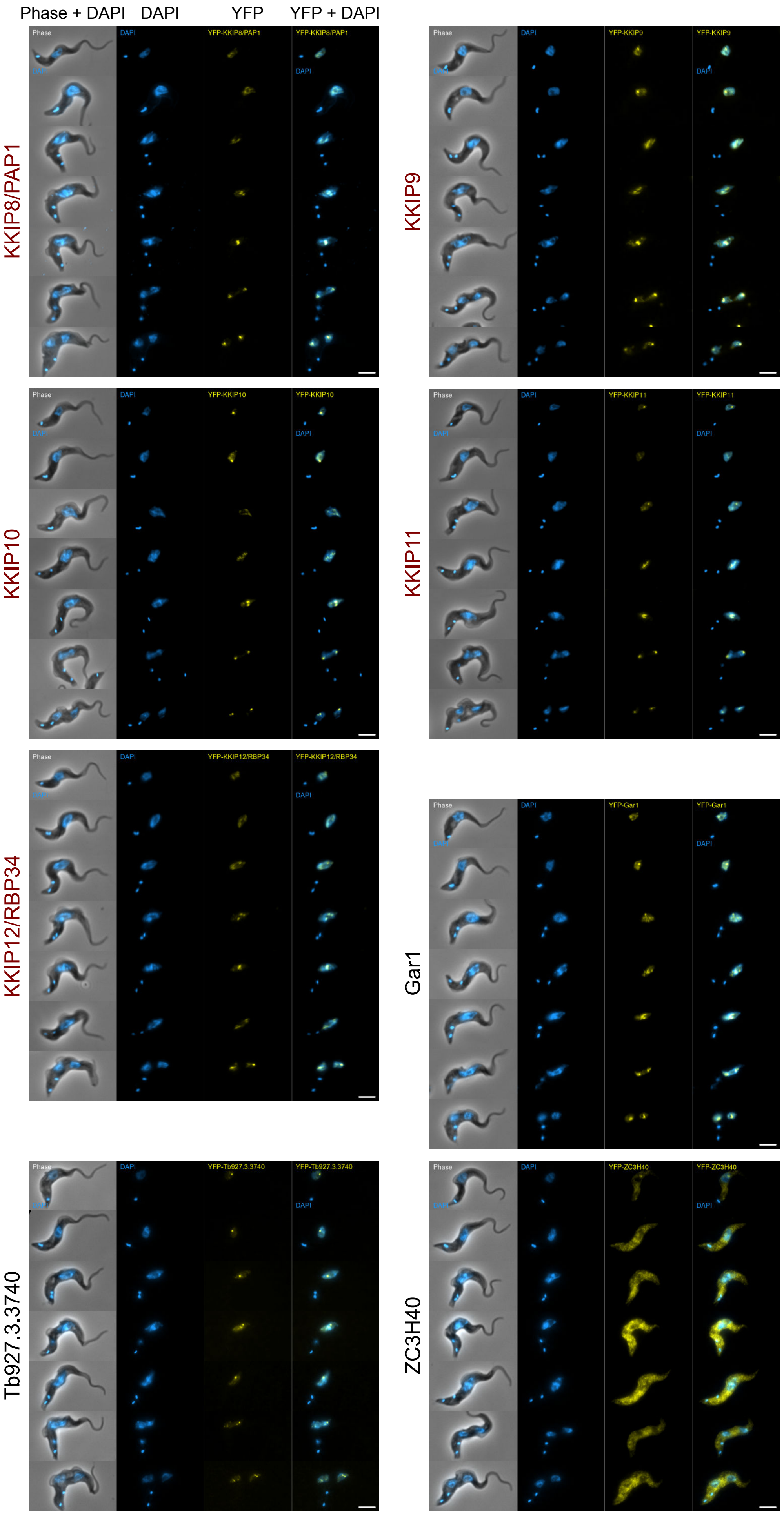

### Supplemental Figure S4

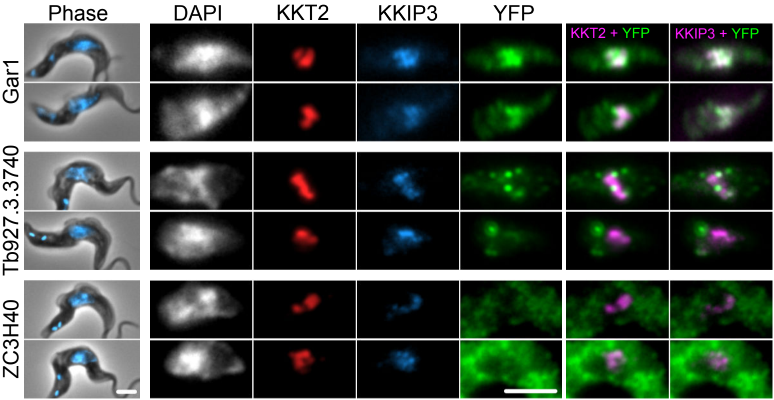

Supplemental Figure S5

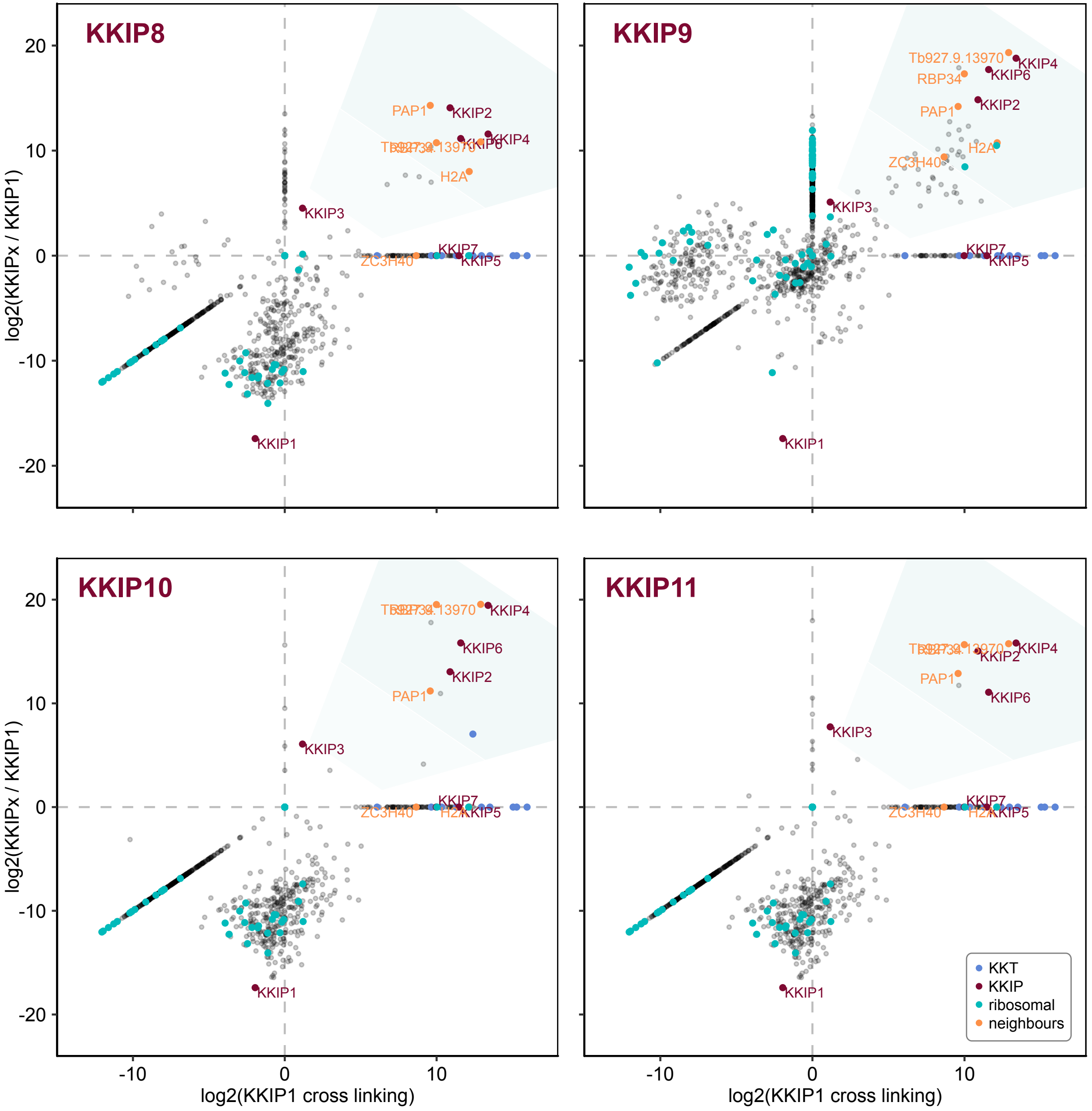

### Supplemental Figure S6

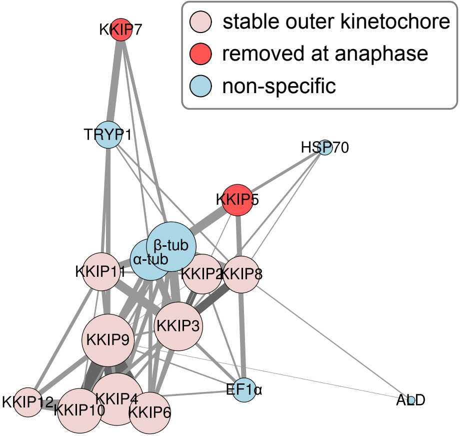
